# AMICI: Attention Mechanism Interpretation of Cell-cell Interactions

**DOI:** 10.1101/2025.09.22.677860

**Authors:** Justin Hong, Khushi Desai, Tu Duyen Nguyen, Achille Nazaret, Nathan Levy, Can Ergen, George Plitas, Elham Azizi

## Abstract

Spatial transcriptomic data enable study of cell–cell communication, yet current analysis tools often fail to provide dynamic, interpretable estimates of interactions and their spatial range across tissue. We present AMICI, an interpretable attention framework that jointly estimates interaction length scales, adaptively resolves sender–receiver subpopulations, and links communication to downstream gene programs. AMICI recovers ground-truth interactions in semi-synthetic data, uncovers gene programs linked to cell communication in the mouse cortex, and reveals length-scale-dependent tumor–immune signaling that reinforces estrogen receptor (ER) programs in breast cancer.

## 1 Main

Cell-cell communication orchestrates a broad spectrum of biological responses, from tissue development and homeostasis to disease progression. Spatial transcriptomics at single-cell resolution offers an unprecedented opportunity to quantify these interactions *in situ* within their native tissue context [1]. Yet, current computational approaches capture only limited aspects of this complex dialogue. Most existing methods primarily rely on the spatial proximity of coarse cell types or aggregate expression patterns associated with known signaling pathways. These include spatial colocalization methods, which identify cell types that co-occur more frequently than at random [2], and interaction models that detect significantly high expression of ligand and receptor gene pairs appearing within close proximity [3, 4]. Despite their utility, these approaches fail to capture the dynamic and context-dependent nature of cell-cell communication at single-cell resolution, overlooking how specific subpopulations of a cell type drive context-dependent phenotypic plasticity in receiver cells through transcriptional shifts.

Recent graph-based approaches such as NCEM [5] and NicheDE [6] aim to bridge this gap by correlating neighborhood cell-type compositions with changes in receiver cell gene expression. Nevertheless, they impose rigid neighborhood definitions, either via a fixed radius or a radial basis function (RBF) kernel, which are challenging to tune and inadequate for interactions operating across diverse length scales. Moreover, their dependence on broad cell-type annotations obscures context-specific subpopulations whose phenotypes are shaped by spatial cues. For example, dichotomous labels such as M1 versus M2 macrophages do not capture the continuum of myeloid states, which transition dynamically under local environmental influences [7]. Similarly, spatially restricted or mixed immune subpopulations within tumors can have opposing prognostic associations with patient outcomes in triple-negative breast cancer patients [8], while hypoxia- and angiogenesis-driven spatial arrangements of regulatory T cells are linked to worse prognosis [9].

To address these limitations, methods such as GITIII [10] and CGCom [11] employ algorithms leveraging attention mechanisms [12] to learn interaction strengths as a function of the neighboring cell’s full transcriptomic profile. However, they frequently yield dense attention maps that are often difficult to interpret and prone to false positives, limiting biological utility.

We propose AMICI (Attention-Mechanism Interpretation of Cell–cell Interactions), which employs an interpretable, attention-based computational framework designed to adaptively learn interactions across varying spatial scales, resolve spatially dependent interacting sub-populations of cells, and link interactions with downstream functional impact in receiver cells. Furthermore, AMICI’s objective function involves sparsity-inducing regularization over the attention scores, ensuring that the model isolates which neighbors, and at what distances, modulate receiver transcriptional programs.

## 2 Results

### The AMICI framework

One key component of cell-cell interaction is the downstream impact of the *sender* cell on the *receiver* cell’s phenotype, which can now be measured with single-cell resolution spatial transcriptomics platforms with expanded panels measuring hundreds to thousands of genes [13, 14, 15]. However, inferring the downstream effect requires identifying the relevant cell subpopulations and the length scales at which they interact, beyond what a cell-type co-localization analysis can provide.

AMICI is a scalable framework that leverages a multi-headed attention module [12] to model each aspect of these interactions in an interpretable manner, but does not directly reuse the attention design from large language models (LLMs). In the context of language modeling, a word *“attends”* to its neighboring context based on the relative position in the sentence as well as the semantic importance of each contextual word. While LLM attention is designed for sequences of words, cells in tissue are not sequential but embedded in 2D or 3D space, where influence is dependent on distance and spatial context. AMICI thus redefines attention for spatial transcriptomics where a receiver cell attends to neighboring sender cells, and the attention-weighted aggregation of their influences determines the receiver cell’s phenotype (**Fig. 1a**). Specifically, during training, AMICI masks the receiver’s own expression profile and learns to reconstruct it from its neighbors.

**Figure 1:**
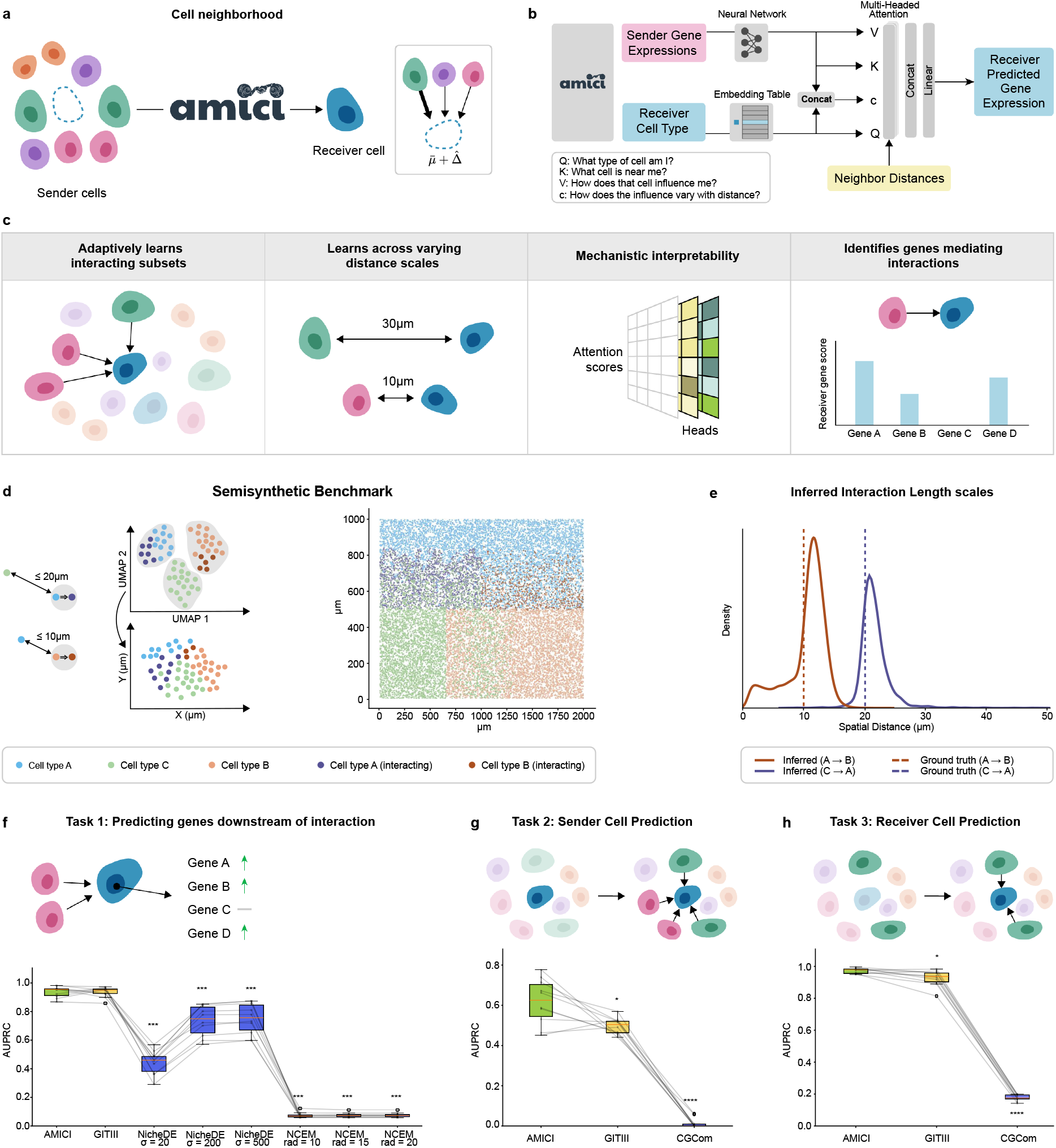
**a**. AMICI simulates the gene expression of a masked receiver cell by reconstructing it from weighted neighbor contributions, capturing shifts from the baseline mean of that cell type. **b**. AMICI model architecture detailing the inputs and prediction outputs to the multi-headed attention layer. A more detailed illustration of the architecture is presented in **Fig. S1. c**. AMICI design choices enable several key advantages over existing methods for inferring cell-cell interactions. **d**. Process of generating the semi-synthetic dataset (**left**) and spatial cell-type distribution of the semi-synthetic dataset (**right**). **e**. Distribution of AMICI’s inferred length scales for an example of the semi-synthetic dataset (solid line) and the corresponding ground truth interaction length scales used to generate the dataset (dashed line). **f**. Area under the precision-recall curve (AUPRC) across 10 technical semi-synthetic dataset replicates for predicting genes downstream of each ground truth interaction for each relevant model. We present NCEM and NicheDE results with varying bandwidth (*σ* or radius (rad)) settings. We denote *p*-value *<* 0.001 as ****, *<* 0.005 as ***, *<* 0.01 as ** and *<* 0.05 as *. Compared to AMICI’s AUPRC scores: NicheDE (*σ* = 20) had *p* = 0.0013, NicheDE (*σ* = 200) had *p* = 0.0013, NicheDE (*σ* = 500) had *p* = 0.0023, NCEM (rad = 10) had *p* = 0.0013, NCEM (rad = 15) had *p* = 0.0013, and NCEM (rad = 20) had *p* = 0.0013. **g**. AUPRC across semi-synthetic dataset replicates for predicting interacting sender cells for the relevant models. GITIII had *p* = 0.028 and CGCom had *p* = 0.0003. **h**. AUPRC across semi-synthetic dataset replicates for predicting interacting receiver cells for the relevant models. GITIII had *p* = 0.028 and CGCom had *p* = 0.0004.

Importantly, the inferred attention weights are a function of both the neighboring cells’ phenotypes, and the distance to these neighbors, enabling AMICI to learn the relevant sender cell populations and the distance at which they influence the receiver cell (**Fig. 1b, Supp. Fig. S1**; **Methods**). By parameterizing attention with a monotonically decreasing function of distance, AMICI can capture interactions across varying spatial scales. With sparsity penalties, AMICI isolates a small number of influential neighbors, mitigating the number of spurious attention scores (**Fig. 1c**). This definition of attention is uniquely suited to this problem as it can adaptively weight heterogeneous neighbors by phenotype and distance, capture multiple length scales through multi-head design, and yield interpretable, biologically grounded communication maps.

Finally, AMICI identifies the downstream transcriptional impact on the receiver cells by using a linear decoder to reconstruct each receiver’s gene expression profile from its neighbors. This architecture ensures a direct, interpretable relationship between the attention scores and the implied influence of each sender cell on the receiver cell. At the population level, AMICI isolates which genes depend on each interaction by evaluating the impact of ablating an entire group of sender cells. With this interpretable framework, AMICI thus moves beyond abstract attention scores to provide concrete, experimentally testable hypotheses that explain how local tissue context reshapes cell states through intercellular interactions.

### Evaluation on Semi-synthetic Data

To evaluate whether AMICI can effectively recover various aspects of cell-cell interactions, we first generated a semi-synthetic spatial dataset with known ground-truth interaction rules. In particular, we constructed a three-cell-type simulation from a single-cell PBMC dataset containing 68, 579 cells and 525 genes [16], with two defined interactions operating at different length scales. For each interaction, we specified a receiver cell type sampled from an “interacting” subpopulation of a cell type only when within the required range of a cell with the corresponding sender type (**Fig. 1d**). In this way, the interacting subpopulation of a cell type carries a distinct set of differentially expressed genes activated only in the presence of the appropriate sender. We assigned the spatial coordinates of the three cell types according to a gradient pattern to generate neighborhoods containing both interacting and non-interacting pairs of cells (**Fig. 1e**; see **Methods**).

AMICI accurately recapitulated the interaction length scales over 10 technical replicates of the semi-synthetic generation process, a feature absent in other tested methods (**Fig. 1f**). This demonstrates that AMICI’s distance-dependent attention mechanism enables the model to learn not only which cells interact, but also how far their influence extends.

We next benchmarked AMICI against several state-of-the-art methods that model per-cell neighborhood effects (NCEM [5], NicheDE [6], GITIII [10], and CGCom [11]), using three metrics designed to capture each method’s ability to recover different aspects of the ground-truth interaction. To evaluate performance comprehensively, we designed three tasks testing whether (1) the models can recover the genes up-regulated downstream of interactions (i.e., ground-truth positively differentially expressed genes in receiver cells), (2) how accurately each method can determine which neighbors act as true senders, and (3) how well they can distinguish which receiver cells were engaged in an interaction. For each task, we computed the area under the precision-recall curve (AUPRC) against the ground-truth values.

On the downstream gene prediction task (**Fig. 1g**) AMICI consistently matched the performance of GITIII and outperformed NicheDE and NCEM even with tuning the model hyperparameters, in particular the bandwidth, reflecting AMICI’s ability to capture significant gene expression contribution towards an interaction. Additionally, AMICI outperformed both GITIII and CGCom on identifying interacting receivers and senders, which demonstrates the importance of regularization in how it mitigates false positive attention scores between cells (**Fig. 1h**, **Fig. S2a-c**). NCEM and NicheDE were not evaluated on the interacting sender and receiver cell prediction tasks, as they can only obtain global predictions at the cell-type level, ignoring their variability across cells and spatial context.

Together, these results demonstrate that AMICI uniquely captures multiple facets of cell–cell communication: which senders and receivers interact, the effective distance of their interactions, and the downstream transcriptional programs involved. Competing methods are limited to subsets of these tasks and fail to recover interaction ranges or identify individual interacting cells, providing limiting mechanistic insights.

### AMICI identifies gene programs shaped by cell communication in the mouse cortex

We next applied AMICI to a spatial transcriptomic MERFISH dataset profiling the mouse primary motor cortex, a region with well-characterized spatial organization and communication patterns. This dataset comprised 64 slices from two mice with 284, 098 segmented cells and 254 genes [2].

Among all cell types, AMICI identified strong interactions from oligodendrocytes and layers 2/3 intratelencephalic (IT) neurons to astrocytes (**Fig. 2a, Supp. Fig. S2d**). Investigating the downstream genes modulated by these interactions, we found *Igfbp5* and *Gfap* to be top genes up-regulated in astrocytes receiving signal from oligodendrocytes. *Gfap* expression in astrocytes is known to play a key role in their interactions with oligodendrocytes during remyelination [17]. Similarly, we found *Cux2* was significantly up-regulated in astrocytes interacting with oligodendrocytes. *Cux2* is known to be involved in the proliferation of cells in the subventricular zone, which includes astrocytes [18] (**Fig. 2b**). These results validate that AMICI can recover meaningful and previously-characterized interactions and provide further interpretation with identifying significant gene sets influenced by the interactions.

**Figure 2:**
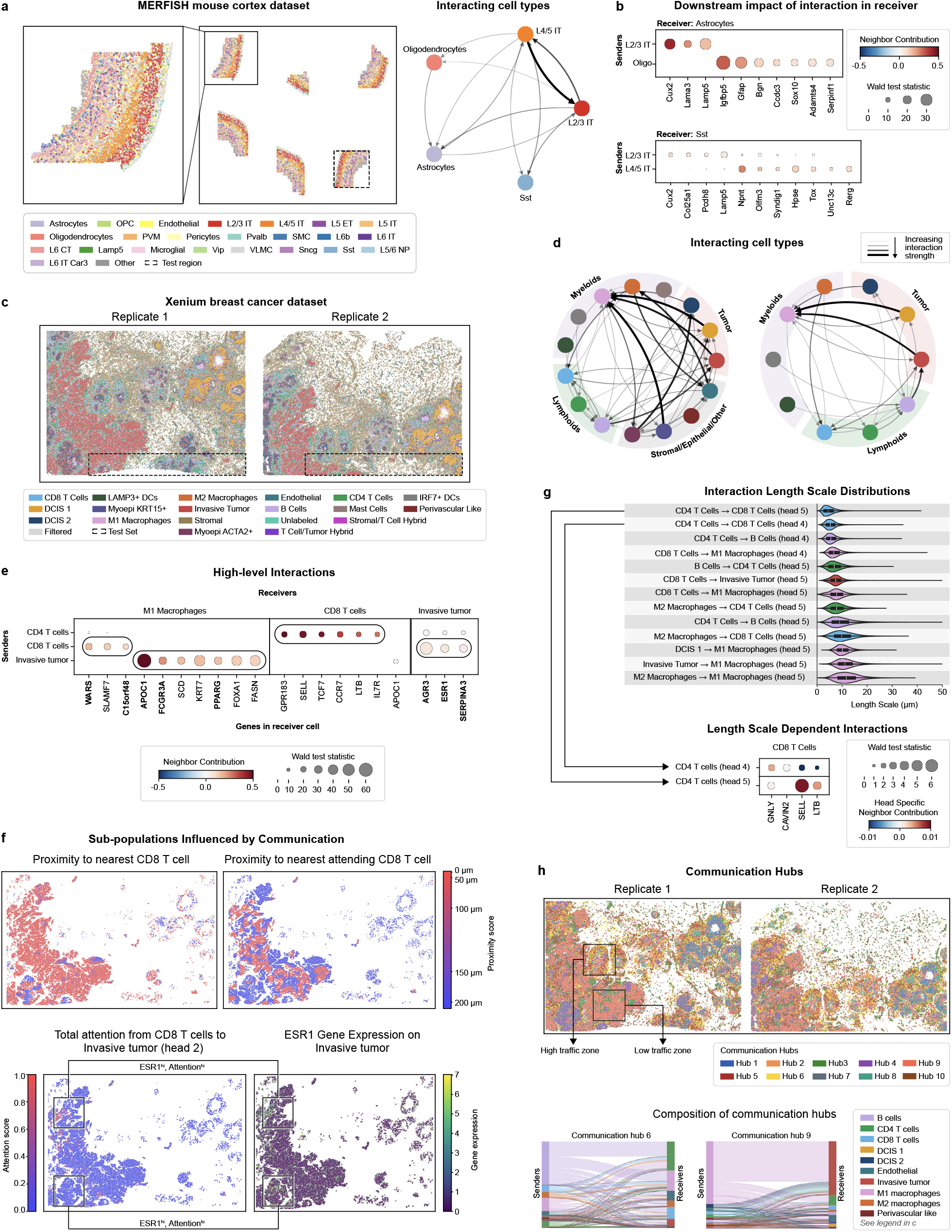
**a**. Spatial cell-type distribution of MERFISH mouse cortex dataset containing 6 samples, with 1 sample held out as the test region (**left**) and directed graph of subset of inferred interacting cell types of interest (**right**). The full graph of all interactions is shown in **Supp. Fig. S2. b**. Downstream genes identified by AMICI for Astrocytes and Sst receiver cells and their corresponding interacting sender cell types. **c**. Spatial cell-type distribution of Xenium breast cancer dataset containing two replicate slides. **d**. Interaction network of all cell types (**left**) and interaction network of immune and tumor cell types of interest (**right**). Line thickness indicates interaction strength between cell type pairs. Legend for cell types is the same as in **c. e**. Downstream genes identified by AMICI for various immune and tumor interactions. Genes that were not flagged to be related to potential segmentation artifacts in the dataset are marked in bold. **f**. Spatial proximity scores for invasive tumor cells to CD8^+^ T cell populations (**top**). High attention from CD8^+^ T cells to invasive tumor overlapping with corresponding high *ESR1* expression at the tumor boundary as a result of interaction, and low attention overlapping with high *ESR1* expression due to external factors in the dataset (**bottom**). **g**. Inferred length scale distributions for various immune and tumor cell-type interactions and corresponding attention head (distribution across all heads is shown in **Supp. Fig. S3c**). Downstream genes influenced by different modes of interactions between the same cell types (CD4 T cell to CD8 T cell) but with varying length scales captured on multiple heads. **h**. Spatial distribution of cells colored by communication hubs identified by AMICI’s inferred signaling patterns, with high-traffic and low-traffic zones (**top**). Communication hubs reflect greater complexity than standard neighborhood niches (**Supp. Fig. S8**). Composition of interacting sender and receiver cell types in communication hubs 3 and 6 (**bottom**) (see all hubs in **Supp. Fig. S8b**).

### AMICI characterizes the landscape of tumor-immune interactions in breast cancer

To evaluate AMICI in a more biologically complex and heterogeneous context, we applied it to a breast cancer dataset collected and profiled using the 10X Genomics Xenium asssay [13]. This dataset composed of two formalin-fixed, paraffin-embedded (FFPE) sections with a total of 286, 532 segmented cells. We first validated that AMICI recovers similar interactions when trained separately on the two slices (**Supp. Fig. S3a-b**). We then jointly fit AMICI to both replicates to increase statistical power for detecting recurrent interactions, withholding contiguous test regions from each slide (**Fig. 2c**). Stromal cells were excluded in training as they were highly susceptible to missegmentation issues, creating spurious interactions with neighboring cells (**Supp. Fig. S4a**).

We examined several high-confidence immune–tumor and immune–immune global interactions identified by AMICI (**Fig. 2c,d, Supp. Fig. S5**). Among the strongest interactions we found CD4^+^ T cell–driven activation of CD8^+^ T cells, marked by upregulation of *TCF7* and *IL7R* [19], and invasive tumor cell influences on M1 macrophages, reflected in elevated *APOC1* expression. This result suggests that CD4^+^ T cell help sustains a reservoir of stem-like, memory-precursor CD8^+^ T cells capable of continuous effector replenishment and enhanced responsiveness to antigenic stimulation and checkpoint blockade [20] [21]. Progenitor TCF7^+^ CD8^+^ T cells are known to co-localize with CD4^+^ helper cells forming tertiary lymphoid structures (TLS) in a cancer tissue microenvironment, which also explains the rise of potential segmentation flags for this interaction [22]. Conversely, given that *APOC1* inhibition reprograms M2 to M1 macrophages via ferroptosis [23], our results also suggest tumor-mediated reinforcement of immunosuppressive macrophage states. Furthermore, invasive tumor cells proximal to CD8^+^ T cells showed upregulated *AGR3, ESR1*, and *SERPINA3. AGR3* promotes tumor proliferation and migration, and *ESR1* is known to regulate *AGR3* in estrogen receptor-positive (ER+) breast cancer [24].

To further analyze this result, we leveraged AMICI’s capability in identifying spatially localized receivers that are influenced by subpopulations of high-interacting senders, moving beyond global cell-type interaction patterns. Interestingly, AMICI’s attention scores revealed spatially distinct invasive tumor subpopulations that are phenotypically impacted by nearby CD8^+^ T cells through *ESR1* expression (**Fig. 2e, Supp. Fig. S6**). This striking result implies that invasive tumor regions may be becoming more ER-dependent as a result of immune interactions beyond known intrinsic tumor cell cycle [25]. This finding is consistent with prior observations that tumor-infiltrating lymphocytes (TILs) in luminal breast cancers are paradoxically associated with worse survival despite predicting chemotherapy response [26, 27]. This phenomenon could be potentially explained by CD8^+^ T cells reinforcing ER signaling through upregulation of *AGR3* and *ESR1*, thereby promoting ER-dependent proliferation and mediating tamoxifen resistance by activating ER signaling despite receptor blockade [24, 28]. At the same time, the upregulation of *SERPINA3* has been reported to reduce resistance of aromatase inhibitors, indicating that understanding tumor-immune interactions can help select more effective pharmacologic targeting of the estrogen pathway in breast cancer [29] (**Fig. 2e**). We additionally evaluated predicted genes on whether they were potentially confounded by segmentation artifacts (**Methods**). A gene was labeled as potentially confounded if it was more expressed in the sender cells than the corresponding receiver cells(**Fig. 2e, Methods, Supp. Fig. S4b**). This analysis confirmed that ER-program genes are not a result of segmentation artifacts. Competing methods were unable to identify genes mediating interactions in the complex tumor tissue, producing only interactions we identified as segmentation-related artifacts (see **Methods**).

By comparing the inferred spatial length scales of these interactions, we found that immune–immune signaling was generally observed at shorter distances than immune–tumor communication (**Fig. 2g**). This result aligns with longer-range tumor-immune communication, which often occurs through cytokine, secreted factor signaling, or hypoxia [30], and the importance of local immune synapse structures for antigen-presenting cells to activate T cell function [31]. We analyzed genes that correspond to varying length scales corresponding to an interaction (learned by different attention heads; see **Methods**). For instance, CD4^+^ T cells induced cytotoxic effector phenotypes in CD8^+^ T cells only through *GNLY* [32] only at a shorter range, while promoting memory CD8^+^ T cell phenotypes through *SELL* and *LTB* [33] occurs at longer ranges as well.

Extending AMICI’s ability to capture functionally relevant subpopulations, we identified broader spatial patterns of communication hubs characterized by types of interactions across the tissue (**Fig. 2h**, **Fig. S7**). These communication hubs capture regions where cells actively influence each other’s phenotypes rather than static niche composition (**Supp. Fig. S8a**). For example, Hub 6 showed diverse immune-immune interactions localized outside invasive tumor boundaries, while Hub 3 was dominated by M1 macrophages influencing tumor cells with minimal other immune involvement, suggesting immune evasion or reduced infiltration. This definition of communication hubs according to diverse signaling patterns reflects a more complex picture than neighborhood compositions (**Supp. Fig. S8a,b**).

## 3 Discussion

Cell-cell interactions follow complex mechanisms that incur cascading effects in a tissue microenvironment. Previous works have attempted to model these interactions using various forms of observable data such as gene expression or co-localization of neighboring cell types. However, these methods present several limitations: they infer interactions based on a fixed radius, whereas cells may interact over different distances; they rely on the resolution of cell-type annotations, which limits the subpopulations of cells from which interactions are inferred; and they are limited in interpretability due to the lack of regularization on attention-based algorithms. To address these issues, we present AMICI, an interpretable and scalable attention-based model, which integrates information from all neighboring cells in 2D or 3D tissue to infer who interacts with whom, at what distance, and which downstream transcriptional programs are modulated.

In semi-synthetic benchmarks, AMICI uniquely recovered all facets of the ground-truth interactions, including sender and receiver identities, interaction length scales, and downstream target genes, demonstrating its capacity to capture complete interaction rules. In the mouse cortex, AMICI identified previously characterized astrocyte–oligodendrocyte signaling programs, validating its ability to uncover biologically established pathways in well-mapped tissues. Applied to breast cancer, AMICI revealed clinically relevant immune-tumor interactions suggesting that ER expression can be influenced by tumor-cell extrinsic factors. While this may offer a possible explanation for the clinical observation that ER+ breast cancers with a dense lymphocytic infiltrate have a worse prognosis, it also highlights a potential therapeutic opportunity, as tumor-immune interactions could sensitize these tumors to aromatase inhibitors or selective ER degraders (SERDs).

AMICI thus consistently captured biologically relevant interactions across datasets with diverse length scales and complexities. We anticipate that future improvements in segmentation algorithms and training on larger cohorts will enable AMICI to improve across broader applications, with implications for deriving fundamental mechanisms and biomarkers for triaging patients and guiding therapy design.

## 4 Methods

### 4.1 Attention-Mechanism-Based Interpretation of Cell-cell Interactions

AMICI is an interpretable attention-mechanism-based model that predicts the gene expression of a cell given its cell-type annotation, its neighbors’ gene expression, and the relative distances to its neighbors. We emphasize that the main takeaway from the model is the downstream interpretation of the learned parameters and intermediate values after training the model. The entire workflow of AMICI can be split into the following steps: (1) data preprocessing, (2) self-supervised training, (3) model evaluation, and (4) downstream interpretation of attention patterns and ablation scores.

### 4.2 Data Preprocessing

AMICI takes in image-based spatial transcriptomics data (e.g., Xenium, MERFISH, CosMX) as input. Using cell segmentation methods, we can obtain single-cell-resolved gene expression measurements with spatial coordinate annotations. When deciding on which segmentation method to use, it is important to consider the level of background in the transcriptomic technology and/or diffusion of transcripts from their original position. These technical artifacts can be mistaken for cell communication events (e.g., if a neighbor’s transcripts leak into the cell boundary of another cell, this may appear to be an influence of the neighbor cell on the transcriptomic state of the receiving cell) and show up as false positives from cell-cell communication inference algorithms. For this reason, we prefer methods that consider approximate transcriptomic profiles [34, 35, 36, 37] over methods that solely use a nucleus/cell membrane stain for image-based segmentation.

After segmentation, the cells must be annotated using marker genes, or annotations can be transferred from existing study annotations, which may be the case when the data is re-segmented. For annotation transfer, we use ResolVI [37], which additionally attempts to correct background gene expression measurements. AMICI effectively models intra-cell-type-annotation variance since the model takes in the cell-type annotation of the target cell. Therefore, the resolution of the cell-type annotations determines the interactions that AMICI can capture. We suggest starting from a high-resolution annotation and merging the annotations based on a prior understanding of the cell types in the dataset. For example, if the annotation resolution separates an exhausted T cell state and a normal circulating T cell state, it would be reasonable to merge these annotations since there is a plausible transition between these states that could be influenced by the neighboring cells. After merging, we suggest filtering out cell-type annotations with particularly low counts. We filter out these cells to avoid learning false-positive interactions that are attributed to overfitting on the few examples present in the dataset. We set this threshold to a minimum of 50 counts per cell in our analyses.

To encourage the model to prioritize the prediction of genes that vary heavily across the cells in the dataset, we filter the genes using the highly variable gene selection method proposed in [38]. The choice of the number of genes to retain is determined roughly by visualizing the histogram of cells with nonzero expression for each gene. After deciding on the ultimate set of genes, we filter out cells with low expression, which is also determined on a per-dataset basis by visual inspection of the count distribution across cells.

Lastly, we normalize the total counts to 1,000 before applying log1p-normalization to the raw count data. We suggest a lower normalization constant of 1,000 due to the generally lower sequencing depth of spatial technologies, but this may be adjusted accordingly based on the technology used.

### 4.3 Self-supervised Training Setup

AMICI is trained in a self-supervised fashion, by masking out the gene expression of cells and passing in the features of neighboring cells as input. To this end, we must first define the neighborhood for any given cell. We denote the target (receiver) cell as *c* ∈ [[1, *N* ]] where *N* denotes the total number of cells in the dataset. We define the neighborhood of *c* as *A*(*c*) ⊂ [[1, *N* ]] *\ c*. Let *M* represent the number of neighbors taken as input for each cell. By default, we choose the 50 nearest neighbors of *c* (i.e., *M* = 50) that do not share the same cell-type annotation as cell *c*. The reason for masking out neighbors of the same cell type is to prevent the model from bypassing neighbor influences by copying the gene expression of the nearest cell in the same annotation group.

For each neighbor *c*^*′*^ in *A*(*c*), we take the observed, log-normalized gene expression levels 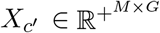, where *G* denotes the number of genes, as well as the relative distances to cell *c*. The distances, *d*(*c, c*^*′*^), are computed as the Euclidean distances between the 2D or 3D spatial coordinates of *c* and each neighbor *c*^*′*^ ∈ *A*(*c*).

The final input to the model is the cell-type annotation of cell *c*, denoted as *t*(*c*) ∈ [[1, *T* ]] where *T* is the number of unique cell-type annotation values.

The objective of the model is to predict the observed, log-normalized gene expression levels of *c*, which we refer to as 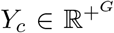. In particular, we aim to minimize the mean-squared error of the predictions *Ŷ* _*c*_ over held-out data.

### 4.4 AMICI Model

The core component of the AMICI model is a multi-headed cross-attention mechanism [12], where the queries are a function of the target cell type, and the keys and values are a function of the neighbors’ gene expression vectors. We modify the attention mechanism to incorporate the influence of distance on the attention score via a monotonically decreasing function class. This choice of function class reflects our assumption that the influence of neighboring cells strictly decreases as they get farther away. The intermediate attention pattern for each head can be summarized as a vector of length *M*, where non-zero attentions correspond to a discernible influence on the target cell’s phenotype. In contrast, fully zero attention will correspond to a prediction solely reliant on the target cell’s label, i.e., cell-type-intrinsic mechanisms. Since AMICI only uses a single attention layer, the learned attention patterns can be directly interpreted as pairwise interactions without compounded effects introduced with additional layers.

We now discuss the model architecture in detail. AMICI learns an embedding matrix 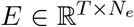 mapping each cell-type annotation, *t*(*c*), to a *N*_*e*_-dimensional vector. In full, the inputs to the attention module are the receiver cell-type embedding (*E*_*c*_, where *c* indexes the columns of *E*), the neighboring cells’ expression values (*{X*_*c*_*′, c*^*′*^ ∈ *A*(*c*)*}*), and the distance between the receiver cell and neighbor cells (*{d*(*c, c*^*′*^), *c*^*′*^ ∈ *A*(*c*)*}*). We assume that the attention patterns should follow some smooth, monotonically decreasing function with respect to the relative distance between the target cell and any given neighbor cell. Rather than using an unconstrained positional embedding (e.g., sinusoidal positional encoding) to incorporate the distances, we construct a function class that expresses our prior assumption.

Additionally, due to the softmax operation, the attention patterns normally sum up to 1 for each attention head. To relax this constraint and allow the heads to drop the attention weight on any neighbor to near zero, we concatenate an “empty” neighbor token to the key vectors and a corresponding “empty” neighbor value vector for this token.

Let **d**_*c*_ := [*d*(*c, A*(*c*)_1_) … *d*(*c, A*(*c*)_*M*_)]. For intermediate values 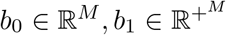, we can express the attention patterns for a select attention head *h* ∈ [[1, *H*]] as:

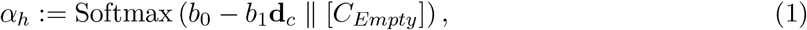

where *C*_*Empty*_ is the pre-softmax weight assigned to the “empty neighbor.” Since *b*_1_ is enforced to be strictly positive, it must be the case that the attention pattern for a given neighbor cell *c*^*′*^ strictly decreases with larger distance values, all other values held fixed.

The neighbor gene expression values are embedded using a multi-layer perceptron (MLP) with a residual connection, 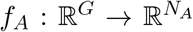. *b*_1_ is computed as a function of the neighbor embeddings, *{f*_*A*_(*X*_*c*_*′*), *c*^*′*^ ∈ *A*(*c*)*}*, and the target cell embedding *E*_*c*_ via neural network, 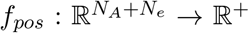, with a final softplus operation to enforce positivity:

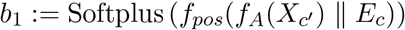

*b*_0_ is computed as an inner product of query and key vectors, similar to in the original transformer architecture [12]. The query vector is computed as a function of the cell-type embedding, whereas the key and value vectors are solely a function of the neighbor gene expression. Each of the vectors is computed with a layer norm applied to the input, followed by a linear layer:

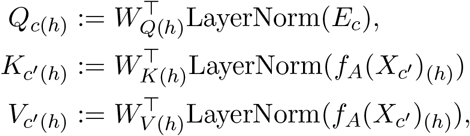

where 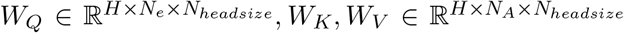 are learned parameters. We can then write *b*_0_ as:

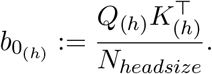

We can plug *b*_0_, *b*_1_ into Equation 1 to compute the attention patterns for each head, *α*_*h*_. The final output of each attention head is then computed as:

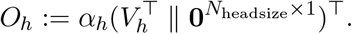

Due to the empty neighbor’s value vectors being a row of zeros, any attention weight placed on the empty neighbor does not contribute to the output of the attention head. The outputs of all the attention heads are then concatenated, flattened, and passed through a linear layer to get an output of dimension *G*, denoted as 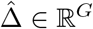. Lastly, we add this output to a vector of the empirical means for each gene for the cell type *t*(*c*), denoted as 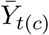. In other words, the model output, 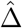, is a predicted residual from the empirical mean of cell type *t*(*c*):

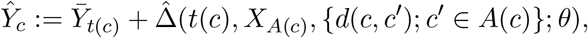

where we denote 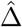 as the AMICI model which outputs the predicted residual from the cell-type mean. The empirical means can also be treated as learnable parameters, which may be necessary in cases where the empirical mean is heavily shifted by interacting cell states. In these cases, fixed empirical means can result in false-positive interactions.

For the attention weights, *α*_*h*_, to be interpretable, we must regularize the MSE loss with a few penalty terms. The first penalty term is a Shannon entropy penalty on the attention weights, *α*_*h*_, which encourages the attention weights to take a sparse pattern if possible. This regularization assumes that a small number of neighbors influence any given cell. The second penalty is an L1 penalty on the value matrix *V*. This penalty encourages the value matrix to take a sparse form and works in conjunction with the entropy penalty to shrink spurious attention weights.

We can write the full loss function for one cell *c* as follows, denoting the penalty coefficients as *λ* with appropriate subscripts:

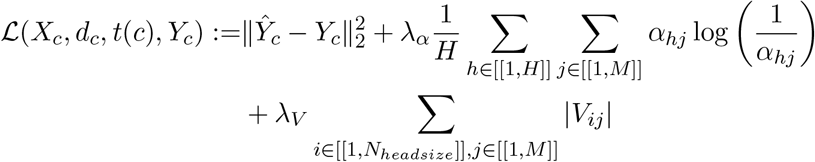

We summarize the key notations employed above in the following table:

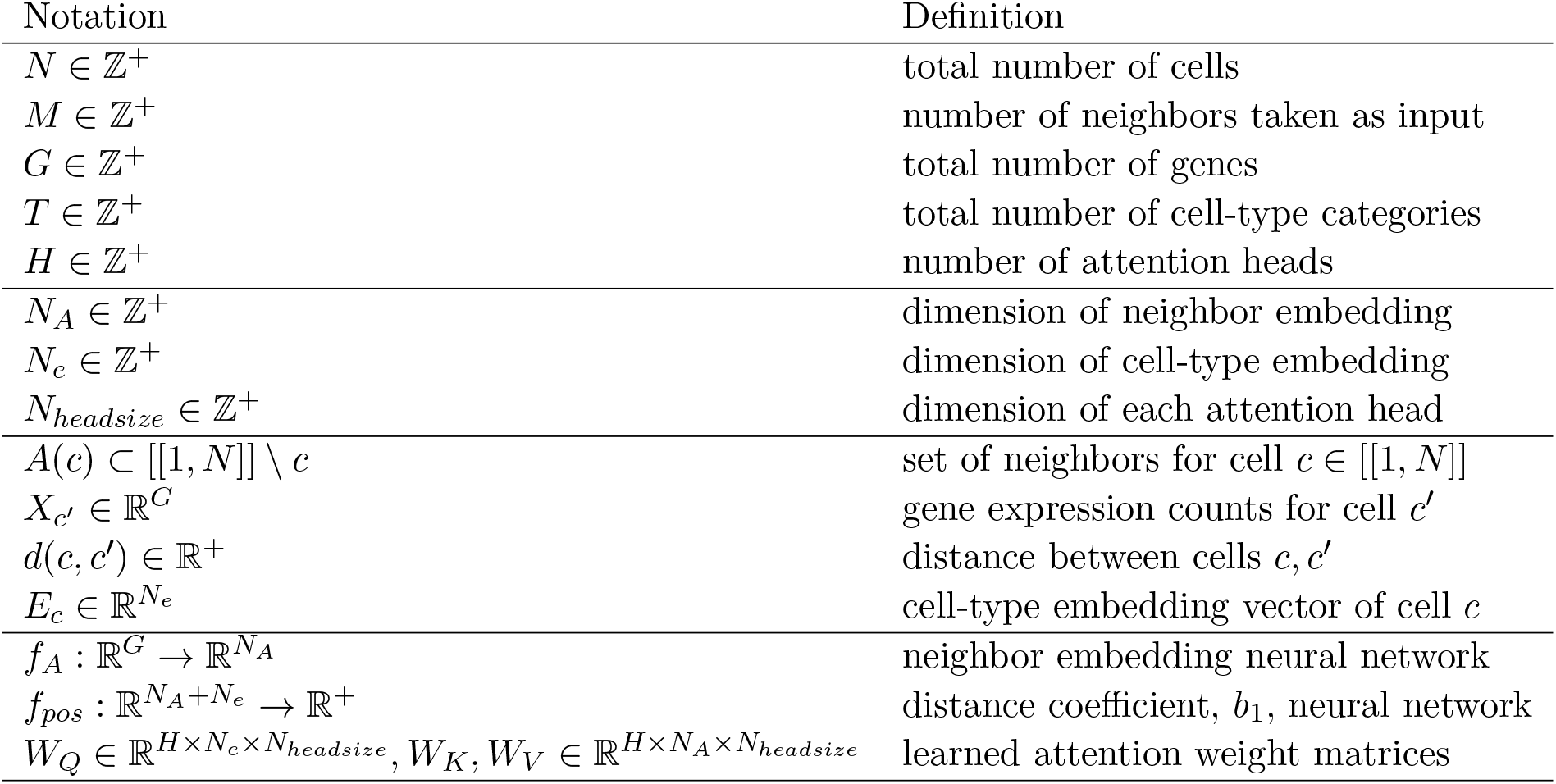

| Notation | Definition |
| --- | --- |
| $N \in \mathbb{Z}^+$ | total number of cells |
| $M \in \mathbb{Z}^+$ | number of neighbors taken as input |
| $G \in \mathbb{Z}^+$ | total number of genes |
| $T \in \mathbb{Z}^+$ | total number of cell-type categories |
| $H \in \mathbb{Z}^+$ | number of attention heads |
| $N_A \in \mathbb{Z}^+$ | dimension of neighbor embedding |
| $N_e \in \mathbb{Z}^+$ | dimension of cell-type embedding |
| $N_{headsize} \in \mathbb{Z}^+$ | dimension of each attention head |
| $A(c) \subset [[1, N]] \setminus c$ | set of neighbors for cell $c \in [[1, N]]$ |
| $X_{c'} \in \mathbb{R}^G$ | gene expression counts for cell $c'$ |
| $d(c, c') \in \mathbb{R}^+$ | distance between cells $c, c'$ |
| $E_c \in \mathbb{R}^{N_e}$ | cell-type embedding vector of cell $c$ |
| $f_A : \mathbb{R}^G \rightarrow \mathbb{R}^{N_A}$ | neighbor embedding neural network |
| $f_{pos} : \mathbb{R}^{N_A+N_e} \rightarrow \mathbb{R}^+$ | distance coefficient, $b_1$, neural network |
| $W_Q \in \mathbb{R}^{H \times N_e \times N_{headsize}}, W_K, W_V \in \mathbb{R}^{H \times N_A \times N_{headsize}}$ | learned attention weight matrices |

### 4.5 Training Procedure

We split the dataset into train, validation, and test splits. The validation set is used for early stopping, which terminates training early if the validation loss does not improve for 10 epochs. We use an Adam optimizer with PyTorch defaults and train for 400 epochs. While the value vector L1 penalty is held fixed throughout training, we increase the attention pattern entropy penalty coefficient, *λ*_*α*_, according to a few hyperparameters. Specifically, we start the coefficient at zero and then increase the value of the penalty linearly to a final value over a range of epochs determined by hyperparameter tuning. For approximately 250, 000 cells AMICI trains for 1 hour using a standard NVIDIA T4 GPU.

### 4.6 Model Evaluation

We use the test mean-squared error to select the hyperparameters for AMICI on any given dataset. In our semi-synthetic benchmarks, we found that the model that minimized the test MSE correctly recovered the ground truth interactions.

For each dataset, we conducted a grid search over the following hyperparameters: `end_attention_penalty, seed, value_l1_penalty_coef, batch_size, lr`.

### 4.7 Model Interpretation

Once the model has been trained, we provide various functions to interpret the learned parameters of the model. Beyond the fixed parameters learned during training, we are interested in the intermediate attention values that vary as a function of the neighboring cells. To this end, we provide a comprehensive set of modules that facilitate both high-level analysis as well as detailed interaction-specific analysis.

#### 4.7.1 Neighbor Ablation for Cell-Type to Cell-Type Influence

To understand the empirical influence of neighboring sender cells on receiver cells, we remove or ablate specific neighbors from the input and compare the predicted gene expression against the unmodified prediction. We define the ablated model predictions as:

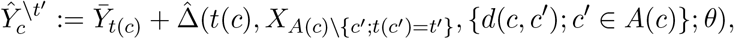

where *t*^*′*^ ∈ [[1, *T* ]] is the ablated cell type. The same ablation process can be done for any arbitrary set of cell indices. With the ablated model predictions, we can understand the contribution of the ablated cell type by computing the difference 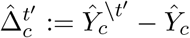.

##### Computing per-gene significance scores

Ultimately, we would like to understand the magnitude of the contributions of a given cell type to the model prediction and whether they are significant with respect to the observed data. As described in [10], we perform a Wald test over a linear model relating the sender cell-type contributions for gene *g*, 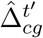, to the observed, normalized gene expression values *X*_*cg*_. This test assumes a standard normal error term. For the null hypothesis that the linear coefficient *β* = 0, the Wald test statistic can be computed as 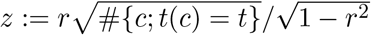 where *r* is the Pearson correlation between 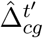 and *X*_*cg*_. The p-value for the two-sided Wald test is then computed as *p* = 2(1 *−* |Φ(*z*)|). The p-values are subsequently adjusted using Benjamini-Hochberg correction.

To summarize the strength of interactions between cell types in a directed fashion, we compute, for every pair of cell types, the number of genes that have a significant adjusted p-value at significance level *α* ≤ 0.05. Note, this interaction strength metric can be biased towards interaction effects well-represented by the gene panel used for the assay. However, this is generally true of any interaction strength metric computed from non-transcriptome-wide ST technologies. We choose to discretize the strength metric to avoid bias towards highly-expressed genes that could dominate a term, such as the sum of sender cell-type contributions.

##### Identification of Relevant Attention Heads

In addition to performing ablation at the full model scale, we can do head-wise neighbor ablations to isolate the contributions of each attention head for a given neighbor cell-type. We define 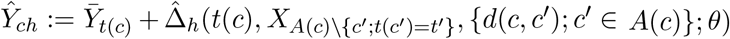 as the prediction made solely with attention head *h*. This ablated prediction is computed by zeroing out the outputs for all attention heads except for head *h*. Then, we can perform the same neighbor ablation as follows: 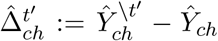. Subsequently, we compute the Wald test statistics and p-values on the 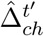 to assess the significance of each head’s contributions.

#### 4.7.2 Counterfactual Attention for Cell-type to Cell-type Interactions

Due to the heterogeneity of cell types in single-cell spatial datasets, we might encounter cell types with disproportionately less cells. As a result, the empirical attention scores may not reflect an equivalent comparison for different cell types across varying distances. We instead use the parameters of the model to compute counterfactual attention scores between two cell types provided the distance between the two cells. Since AMICI learns *Q*_*h*_, *K*_*h*_, and the coefficient *b*_1_, we can compute the attention score given a counterfactual distance between the two cells, *d*^*cf*^. We define the counterfactual attention patterns as: 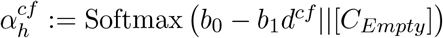

##### Identification of the Length Scales of Interaction

Using AMICI’s learned parameters we can also identify the length scale of an interaction between two cell types. Since the attention patterns follow a monotonically decreasing function, we can take the distance at which the counterfactual attention scores drop to a low threshold, *α*_*h,th*_ as the length scale of that interaction. Reformulating our counterfactual attention pattern definition the length scale distance, we get 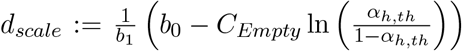. Empirically, we found that *αh,th* = 0.1 works well. It is possible to obtain spurious high counterfactual attention scores for cell types with extremely small distances that are out of distribution for the model’s learned parameters. These may result in a length scale with a very small value despite having negligible empirical attention between the cell types. We apply a filtering procedure to eliminate these false length scales. For a given sender cell type and attention head, we compute the proportion of receivers that have at least one of the sender cell type within its nearest neighbors, at a distance within the median length scale across senders. If this proportion is below a sample threshold, we set the length scale to zero.

#### 4.7.3 Identification of Interacting Receiver Subpopulations

Cell type annotations in single-cell datasets may not accurately reflect the correct resolution at which cell-cell interactions occur. For example, there may be phenotypically distinct subpopulations of tumor cells interacting with immune cells in different spatial regions of the tissue. For the head that explains the most variance for the receiver cell-type of interest, we can accumulate the attention paid to the corresponding neighbor cell type. We can plot these to further visualize how the most influenced receiver cells vary spatially across the tissue, as well as how they vary with respect to their high-attention senders.

#### 4.7.4 Identification of Spatial Communication Hubs

Finding regions of the tissue which contain diverse immune-immune interactions or high tumor activity can help characterize prognostic outcomes and identify target regions where interventions could modulate the tumor-immune interface. We define regions that contain strong interactions with similar neighbor cell types in a microenvironment. For each receiver cell type, we define high-interacting cells as those with the top 10% of nonzero attention scores across all neighboring cells. Then for each cell we construct a cell-type composition vector, where we only count high interacting cells from the neighborhood of the cell. This composition vector is used to build clusters of cells using KMeans with *k* clusters, which we define as communication hubs. The number of hubs (clusters) is determined by optimizing for the best silhouette score over a range of valid values of *k* (**Supp. Fig. S7a**). The silhouette score is computed for every cell *i*,

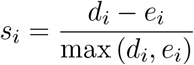

where *d*_*i*_ is the distance from cell *i* to its nearest cluster, and *e*_*i*_ is the average distance from cell *i* to other cells within its own cluster. We computed the adjusted rand index (ARI) and the adjusted mutual information (AMI) scores between regular cell-type composition of the neighborhood and simply the cell type labels to show these clusters are unique to AMICI (**Supp. Fig**. S8).

### 4.8 Semi-synthetic Benchmark

We simulate a semi-synthetic dataset to validate AMICI with ground truth interactions, while utilizing aspects of real single-cell data. We simulate interactions within the dataset as phenotypic shifts in a cell based on the neighboring cells and its distance to those cells. Different cell-types are simulated to interact at varying length scales as this reflects how cell-cell interactions may appear in real datasets.

We expect AMICI to capture these effects as the model learns to predict a shift in the mean expression of a cell using the information its nearest neighbors. By learning a coefficient that varies per neighboring cell, AMICI can identify the different ranges at which interactions occur between one pair of cell types versus another. On the other hand, methods like NCEM and NicheDE may not be able to capture these interactions. NCEM and NicheDE model interactions by neighbor cell-type abundance, which may lead to false positive predictions for cell-type pairs that are not interacting. Furthermore, they use a fixed radius to define their niche and thus cannot determine the varying length scales at which the ground truth interactions occur.

#### 4.8.1 Dataset Generation

We use a single-cell dataset of fresh PBMCs from a healthy donor (Donor A) with 68,000 detected cells [16] as our reference. The dataset is pre-processed by normalizing the gene expression to median total counts, applying log1p-normalization, and filtering for the top 500 highly variable genes.

We train an scVI model on the dataset to generate embeddings for each cell [39]. We then use Leiden clustering detailed in [40] on the scVI embeddings of the cells to create clusters, selecting three of them as our main cell types for the simulation: A, B and C. Our simulation involves two interactions where Cell Type A is the sender for a Cell Type B receiver at a length scale of 10 micrometers, and Cell Type C is the sender for a Cell Type A receiver cell at a length scale of 20 micrometers. We subcluster clusters for Cell Type B and A to create two subclusters for each cell type. One cluster represents the interacting phenotype of the cell type, and another represents a non-interacting state. We create a uniform grid within a rectangle and assign cell types based to different quadrants as their own neighborhoods, with gradient transitions at the boundaries where cell types intersect. The gradient effect is created by using the spatial coordinate’s distance to the dividing quadrant border to determine the probability of assigning that location to its quadrant’s dominant cell type versus the neighboring quadrant’s cell type.

Next, we assign these spatial locations to actual cells sampled from different clusters in our reference dataset. For each receiver cell, we sample from the cell type’s interacting cluster if the corresponding sender is spatially located at a distance within the defined length scale. If there is no sender within that distance for the receiver or the cell type is not a receiver in any interaction, we sample from the non-interacting cluster.

The data generation process includes various stochastic processes, including training scVI, randomly selecting a subset of genes for simulating the interaction, randomly generating uniform spatial locations, and sampling cells from populations of interacting or non-interacting cell clusters. To ensure that AMICI is robust regardless of the stochastic variables in our simulation, we run the dataset generation process on 10 different scVI seeds and assess its performance across all of them.

#### 4.8.2 Training Setup

We split the dataset into a train and a test set. For the test set, we choose a slice of the dataset between the spatial locations of cells where *X*_*c*_ *>* 900 and *X*_*c*_ *<* 1100 to ensure the test set contains all varying spatial neighborhoods. We train AMICI by conducting a sweep over the seed parameter and keep the model with the lowest MSE loss. We find that `value_l1_penalty_coef = 0.00001, end_attention_penalty = 0.1, lr = 0.001, batch_size = 128` works well across all semi-synthetic datasets.

#### 4.8.3 Baseline Models

We benchmark AMICI against four baseline models: NCEM, NicheDE, GITIII and CGCom. Since NCEM and NicheDE use a fixed radius we evaluate their performance on the semi-synthetic simulations with varying niche sizes. We test NCEM on radius sizes 10, 15 and 20, whereas for NicheDE we evaluate on larger niche sizes as well such as 200 and 500, as suggested in the paper. We train GITIII, NCEM and CGCom with their default parameter settings. We run these baseline models on all 10 replicates of the semi-synthetic dataset as well to create an equivalent benchmark to compare AMICI against.

#### 4.8.4 Metrics

We measure the performance of AMICI against other methods on three different tasks: DE gene prediction, interacting sender cell prediction, and interacting receiver cell prediction. We compare the AUPRC of PR curves generated from each baseline model’s predictions on a particular task with that of AMICI. We perform a Mann-Whitney U test to obtain a significance metric (p-values adjusted using Benjamini-Hochberg correction) between AMICI’s AUPRC scores and those of each of the compared baseline methods.

##### DE gene prediction

The ground truth genes are the DE genes between the interacting subtype and the neutral subtype for a given receiver cell type in an interaction, with positive log-fold change scores and p-value *≤* 0.05. We use the Wald-test statistic generated from computing per-gene significance scores as AMICI’s predictor value for this prediction task.

##### Interacting sender cell prediction

For a given interaction with a sender and receiver cell type, a neighbor cell is an interacting sender cell if it is of the sender cell type and is within the length scale of the correct corresponding receiver cell type. Since GITIII uses a different method and may use a different number of nearest neighbors than AMICI, we only assess the model’s ability to predict neighbors that overlap across both models. To predict interacting sender cells, we use the maximum value of AMICI’s empirical attention patterns across all receivers and attention heads for each sender cell.

##### Interacting receiver cell prediction

During the dataset generation, each receiver is sampled as an interacting subtype if a sender cell type is within the distance range of the corresponding sender cell type for that interaction. We use these defined interacting subtypes as the ground truth for the interacting receiver classification. To predict the interacting receiver cells, we use the maximum attention pattern across all senders and attention heads for a particular receiver cell.

### 4.9 Mouse Cortex MERFISH Dataset

The MERFISH dataset contains 64 slices of the mouse brain motor cortex from two different mice. It is composed of 284,098 segmented cells, 254 genes and 24 annotated cell types across the slices. We select mouse 1 sample 4 which contains 6 slices and 33,831 total cells.

#### 4.9.1 Preprocessing

The dataset provides segmented cell boundaries, cell-type labels, and raw gene counts for each cell. We compute the centroids for every cell by taking the average of the *x* and *y* coordinates of the cell boundaries. We normalize the counts to 1,000 and apply log1p-normalization to the data.

#### 4.9.2 Model Training/Selection

We split the dataset into train and test by holding out slice 180 of the 6 slices from the sample as a test set to evaluate AMICI’s performance. We perform a sweep over the parameters, `end_attention_penalty, attention_penalty_schedule, seed, batch_size` and `value_l1_penalty_coef`, taking the model with the lowest test loss.

#### 4.9.3 Benchmarking

We compared against the results reported for the same dataset in the corresponding publications of NCEM [5] and GITIII [10]. Both of these models both recovered similar interactions and associated genes to AMICI.

### 4.10 Breast Cancer Xenium Dataset

The Xenium dataset is comprised of two consecutive 0.4cm^2^ slides of a sample of breast cancer tissue, annotated by a pathologist. Replicate 1 contains 167,780 segmented cells and replicate 2 contains 118,752 segmented cells.

#### 4.10.1 Preprocessing

Some annotated cell types in the dataset were phenotypically quite similar and may incur dominating false positive interactions if they are treated as separate cell types during the training process. As a result we group specifically Proliferating Invasive Tumor and Invasive Tumor together. We additionally noticed that some DCIS 1 and DCIS 2 cells were mis-annotated and lay in incorrect pathologist annotated regions of the slide. To correct for this, we used regions demarcated by the pathologist to assign a boundary between the DCIS 1 and DCIS 2 cells and re-annotated them according to their corresponding regions. We also removed unlabeled or hybrid cell with multiple cell-type labels since the varying phenotypes of these cells may also conflate interactions inferred by AMICI. Finally, since Stromal cells were found to be susceptible to mis-segmentation issues, we filtered out these cells as well.

We use Proseg and ResolVI annotation transfer to re-segment and re-label both replicates of the dataset. We used default parameters for Proseg and assigned cell-type labels based on the probabilities output by ResolVI. To avoid having cells that may be mislabeled due to uncertainty of the annotation algorithm, we removed cells that were labeled with a probability below a certain threshold. We found 0.5 to be a good balance between ensuring certainty of cell-type labels and filtering out too many cells. We also filter out cells with counts below 50 and filter for the top 250 genes, before normalizing to a total of 10,000 counts and applying log1p-normalization. After filtering out cells we are left with a total of 199,912 cells across both datasets.

#### 4.10.2 Model Training/Selection

We split the dataset into train and test by taking a slice of each replicate as the test set on which we evaluate AMICI. We perform a sweep over attention penalty schedule, end attention penalty, seed, and value l1 penalty coef, and keep the model with the lowest MSE loss when evaluated on the held-out test set.

We trained AMICI separately on the two biological replicates and compared the identified interactions against each other. The interactions identified and corresponding significant genes involved in the interactions were consistent across both replicates (see **Supp. Fig. S3a-b**), implying that AMICI generalizes well and demonstrates reproducible interpretability across similar datasets.

#### 4.10.3 Significance Test for Segmentation Effects

Despite re-segmentation with Proseg [36] and ResolVI [37], it is possible to encounter segmentation artifacts in single-cell spatial data between other cell types. We thus design a test to rule out segmentation error impacting prediction of downstream genes in receiver cells. Specifically, we investigate if the expression for a gene *g*, identified by AMICI for a sender-receiver interaction, is significantly higher on the receiver while it is interacting versus on the sender while it is distant from the interaction. If we see that it is not significantly higher on the interacting receiver, we can attribute that expression to segmentation artifacts from the corresponding sender. For a given receiver-sender pair, we define the set of interacting receivers, *r*_*int*_ as the set of cells of the receiver cell-type that are within 20*µ*m from a sender cell. We also define a set of distant senders, *s*_*dist*_ as the set of cells of the sender cell-type that are at least 100*µ*m from a receiver cell. We obtain 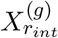 the distribution of gene expression for gene *g* of *r*_*int*_ and 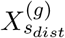. We perform a Mann-Whitney U test where,

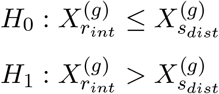

We reject the null hypothesis with resulting *p*-value *<* 0.05.

We note that if a gene can be expressed on both the sender cell type and receiver cell type, this test cannot confidently rule out the impact of mis-segmentation. Thus is it more applicable to marker genes exclusive to cell types.

#### 4.10.4 Benchmarking

We compared against the results reported for the same dataset in the GITIII publication [10]. GITIII recovers a subset of similar genes to AMICI for most interactions; however, these genes are often associated with segmentation errors that we flag and exclude from our downstream analyses.

## Data Availability

All data used for experiments in this work are publicly available. The semi-synthetic experiment was constructed using data from [16], which can be found at https://www.10xgenomics.com/datasets/fresh-68-k-pbm-cs-donor-a-1-standard-1-1-0.

The mouse cortex MERFISH data from [2] can be downloaded from https://doi.brainimagelibrary.org/doi/10.35077/g.21. The breast cancer Xenium data from [13] can be downloaded directly from the 10x Genomics website at https://www.10xgenomics.com/products/xenium-in-situ/preview-dataset-human-breast.

## Code Availability

All code, including source code, dependencies, and scripts to run data analyses featured in this paper, can be found in the GitHub repository: https://github.com/azizilab/amici.

## Acknowledgements

We thank Adam Gayoso, Lingting Shi, Mingxuan Zhang, Yinuo Jin, and Nicholas Beltran for helpful discussions and feedback that greatly improved our work. We would also like to thank Linyue Fan for creating the logo.

This work was supported by the NIH NCI grant R00CA230195, NIH NHGRI grants R01HG012875, R21HG012639, NSF CBET 2144542, and grant number 2022-253560 from the Chan Zuckerberg Initiative DAF, an advised fund of Silicon Valley Community Foundation.

## Author Contributions

J.H. and K.D. contributed equally to this work. J.H. and E.A. conceived the original research questions and developed the conceptual framework. T.D.N. and A.N. explored the initial problem of this project and conducted analyses that helped inform the final approach. J.H. designed the methodology and implemented the first iterations of the method. J.H. designed the semisynthetic benchmark, and J.H. and K.D. implemented the semisynthetic benchmark analyses together. J.H. and K.D. designed and developed the downstream interpretation methods. N.L. and C.E. resegmented and pre-processed the Xenium breast cancer data used in this study. K.D. led and implemented the real dataset analyses. G.P. contributed to the interpretation of breast cancer results and helped edit the paper accordingly. E.A. oversaw the entire project. J.H., K.D., and E.A. wrote the manuscript.

## Supplementary information

### Supplementary Figures

**Figure S1:**
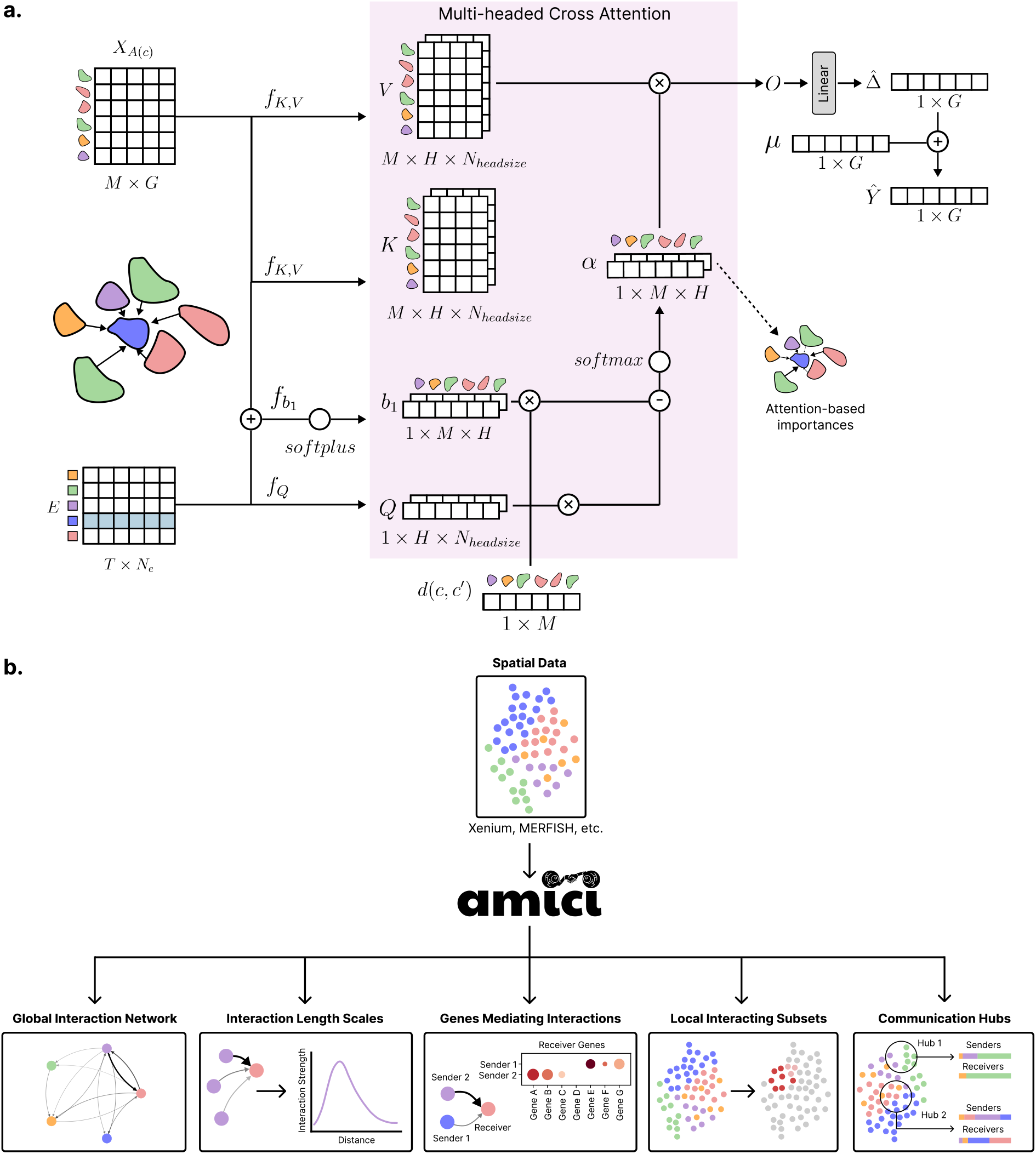
**a**. Multi-headed attention architecture for the masked gene expression prediction task through which AMICI learns its attention scores (see **Methods**). *T* is the number of cell types, *M* represents the number of nearest neighbors for a receiver cell, *H* is the number of heads in the attention model, and *G* is the number of genes in the dataset. **b**. AMICI framework for predicting cell-cell interactions from spatial single-cell data. AMICI’s downstream tasks include: (1) global interaction network prediction showing directed relationships and interaction strengths between cell types; (2) interaction length scale analysis determining the spatial distances at which cell types communicate; (3) gene prediction identifying key genes modulated in receiver cells with specific sender-receiver interactions; (4) local neighborhood analysis identifying receiver cell subsets highly influenced by surrounding cells; (5) communication hub detection revealing distinct interaction patterns and corresponding sender-receiver compositions across tissue regions.

**Figure S2:**
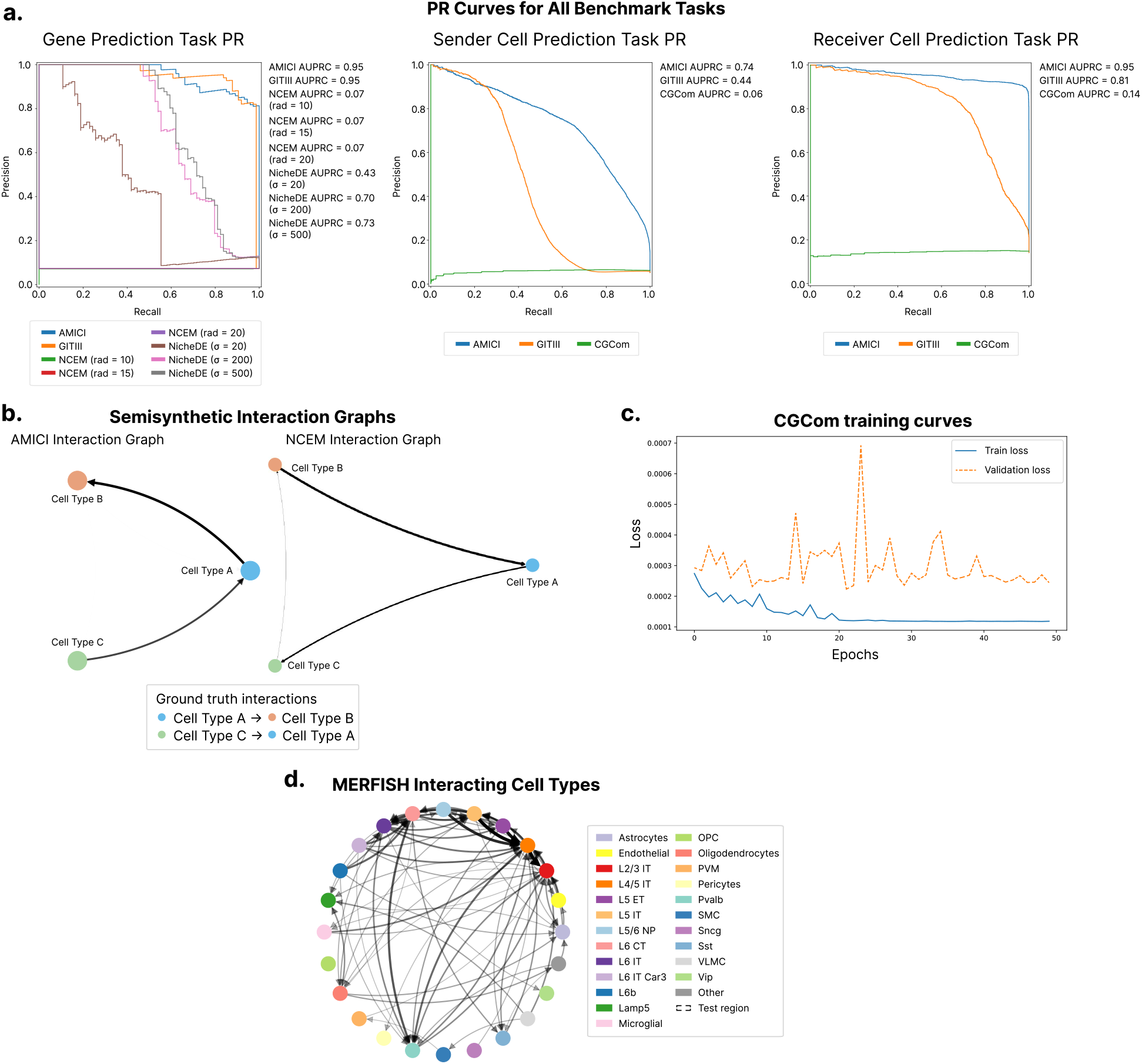
**a**. Precision-recall curves for all relevant models for the gene prediction task (**left**), sender cell prediction task (**center**) and the receiver cell prediction task (**right**) on the semi-synthetic dataset. AMICI outperforms GITIII on the sender and receiver cell prediction tasks as GITIII does not employ regularization, which makes it prone to false positive predictions. **b**. Predicted interaction networks compared to the ground truth interactions in legend. AMICI (**left**) correctly identified interaction direction and relative strength. NCEM (**right**) shows reverse interaction direction and false positive predictions, which accounts for its poor performance in downstream gene prediction tasks. **c**. CGCom training and validation loss curves suggesting overfitting, which expalins lower performance on the sender and receiver cell prediction tasks. **d**. Comprehensive directed interaction network of cell types in the MERFISH mouse cortex dataset. Edge thickness indicates interaction strength between cell type pairs.

**Figure S3:**
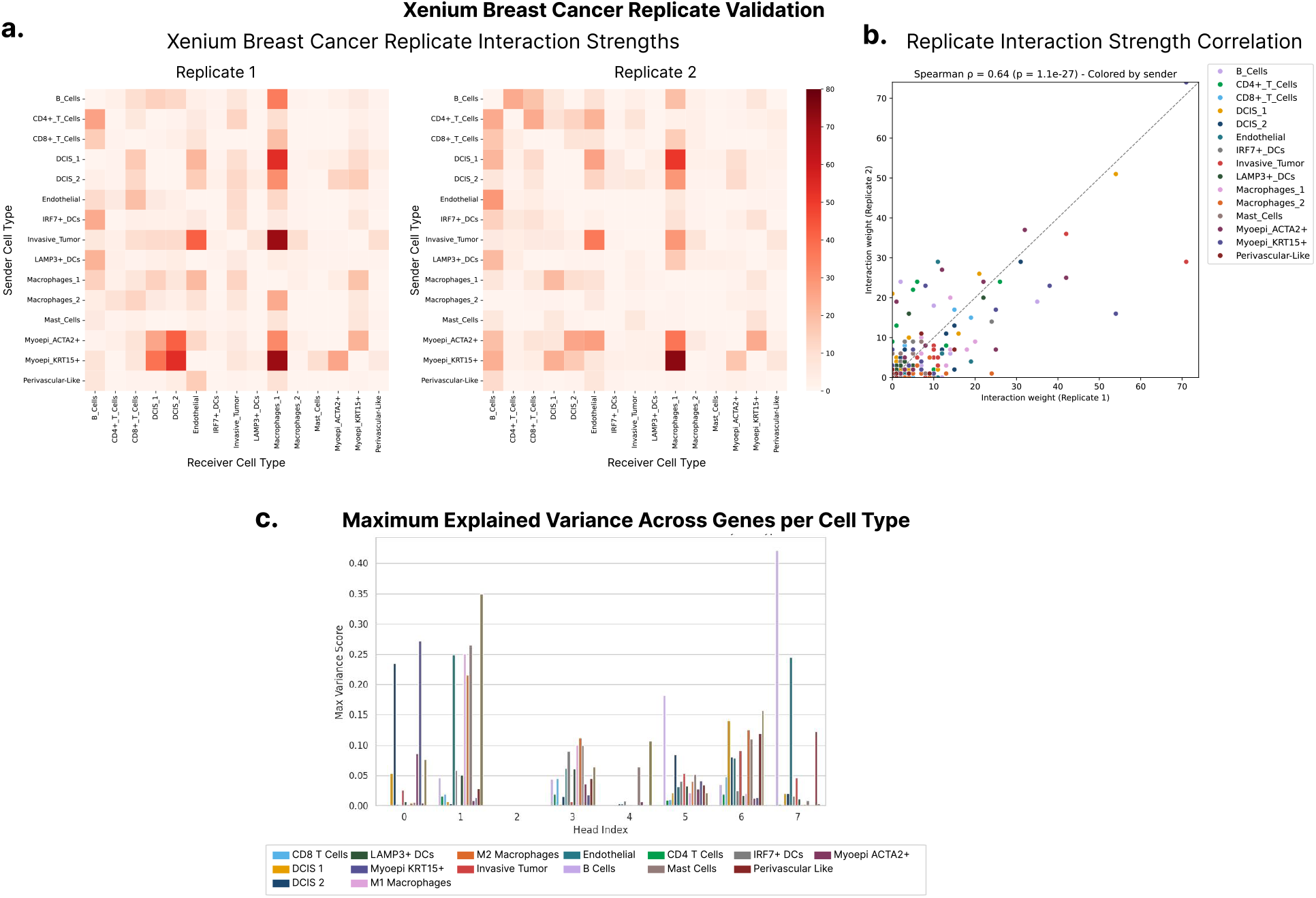
Validation for AMICI’s training and results on two consecutive slides of Xenium breast cancer dataset. **a**. The interaction strength matrix predicted by AMICI between pairs of cell types for replicate 1 (**left**) and replicate 2 (**right**), showing reproducible interaction patterns across the two replicates. **b**. Scatterplot showing high overall correlation for AMICI’s predicted interaction strength scores between replicates. Each dot corresponding to a pair of cell types, colored by the sender cell type. **c**. Maximum explained variance score between predicted gene expression with and without the given head, for each corresponding cell type. The explained variance is spread across multiple heads for different cell types, which confirms the need for multiple attention heads to capture the diverse interactions in complex tissues such as the tumor.

**Figure S4:**
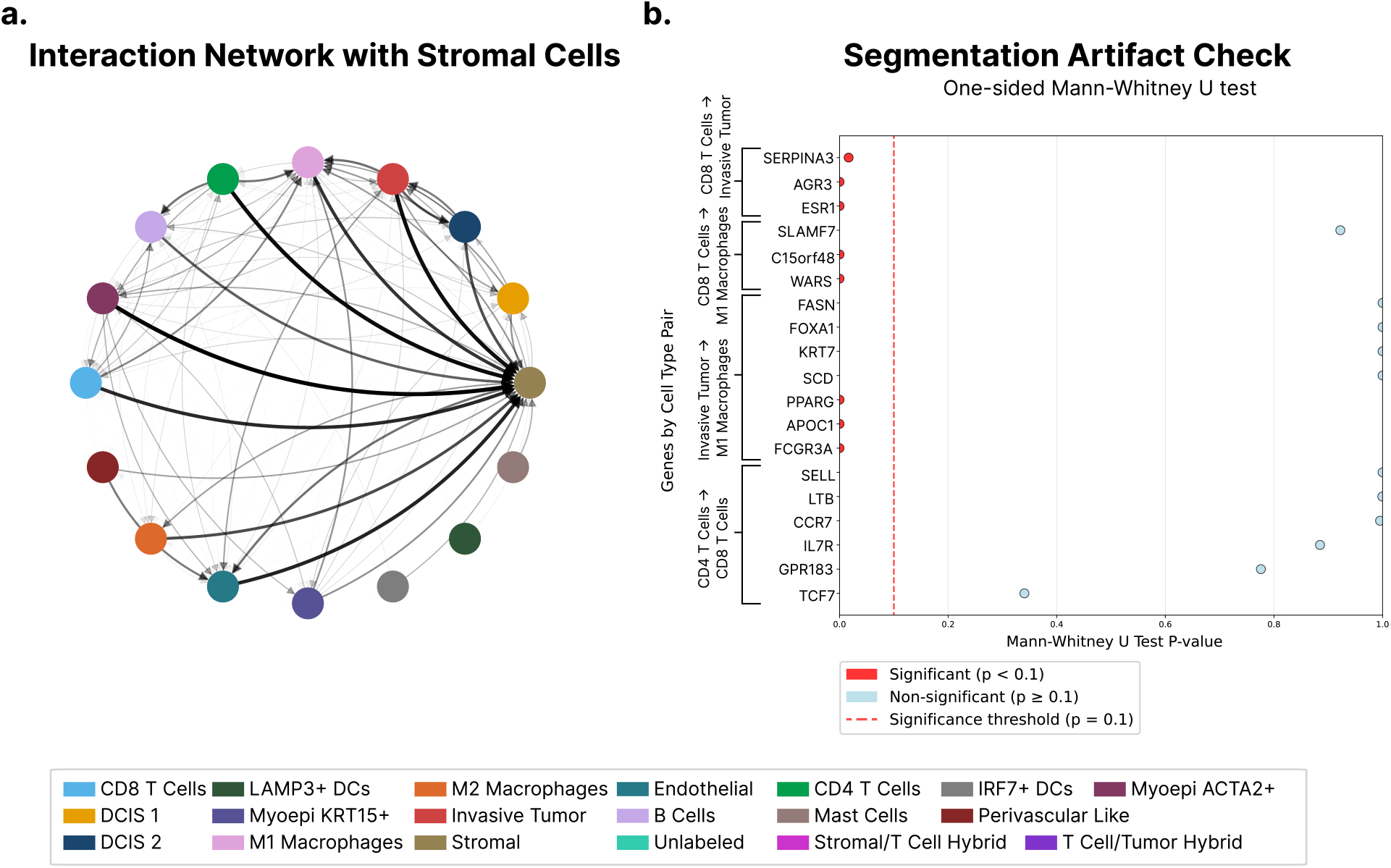
**a**. Directed interaction network of cell types in the Xenium breast cancer dataset when trained on all cell types, including stromal cells. The interaction graph shows strong connections between almost every cell type with the stromal cells as they are prone to segmentation artifacts that arise in Xenium data. The artificially strong stromal interactions skew the relative contributions between other cell types, potentially masking biologically relevant communication patterns. We thus recommend excluding cell types with major segmentation issues dominating the entire tissue. **b**. Statistical validation of genes identified by AMICI using one-sided Mann-Whitney U test. Test evaluates whether genes are significantly higher in receivers near senders versus those farther away, distinguishing true interaction-mediated expression from misaligned transcript artifacts. Red markers indicate validated high-confidence interaction genes as they are unlikely to be influenced by segmentation errors; blue markers show genes that may be due to segmentation errors or alternatively marker expression shared between sender and receiver cell type.

**Figure S5:**
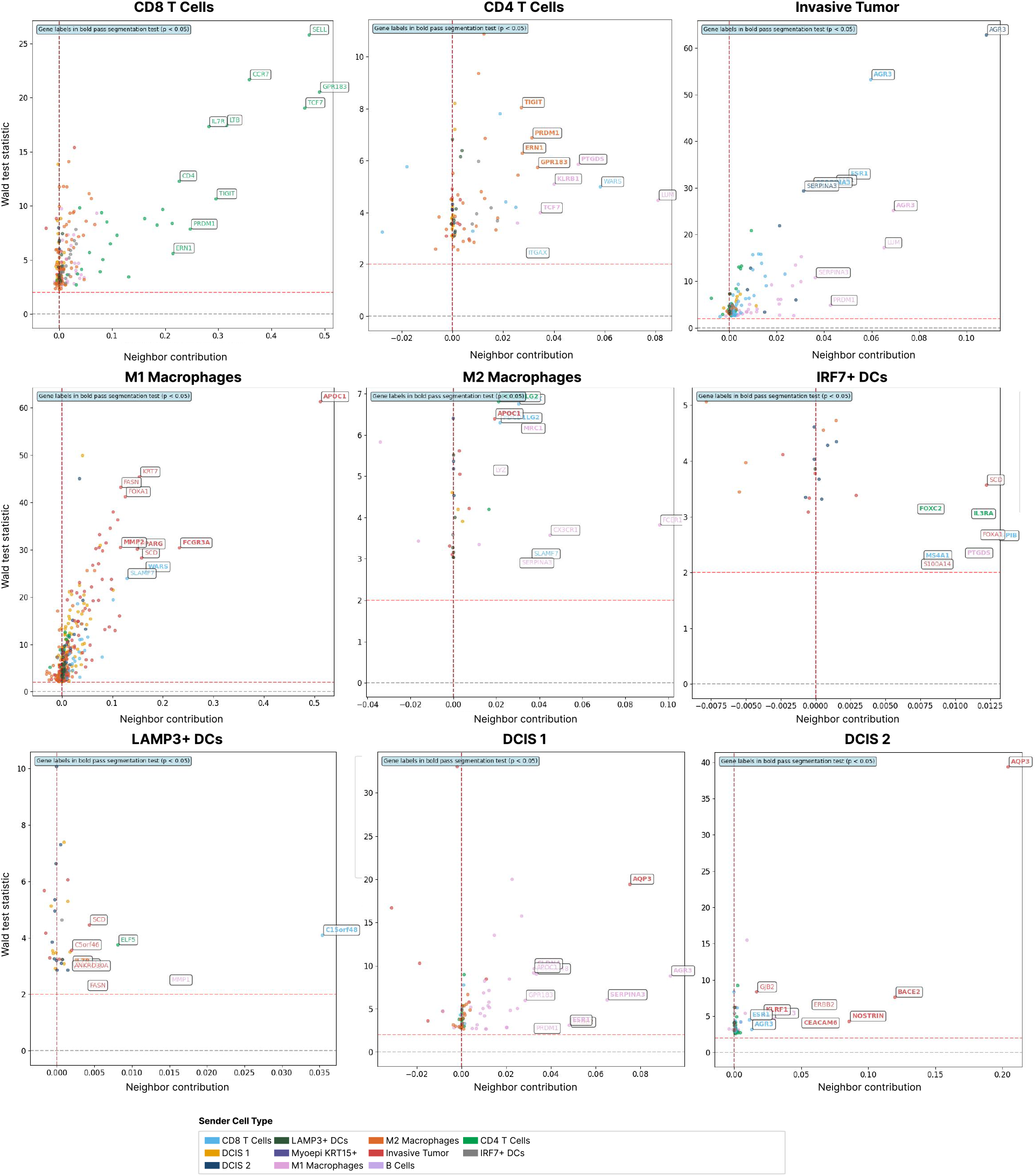
Volcano plot displaying all genes in each receiver cell type and how they are modulated by immune-tumor interactions. Each subplot corresponds to one receiver cell type. Each dot is a gene, colored by influencing sender cell type. The x-axis shows the neighbor contribution score and the y-axis shows the Wald test statistic for each gene. Top 10 genes sorted by neighbor contribution have been labeled and bolded labels highlight genes that were not flagged as potential segmentation artifacts.

**Figure S6:**
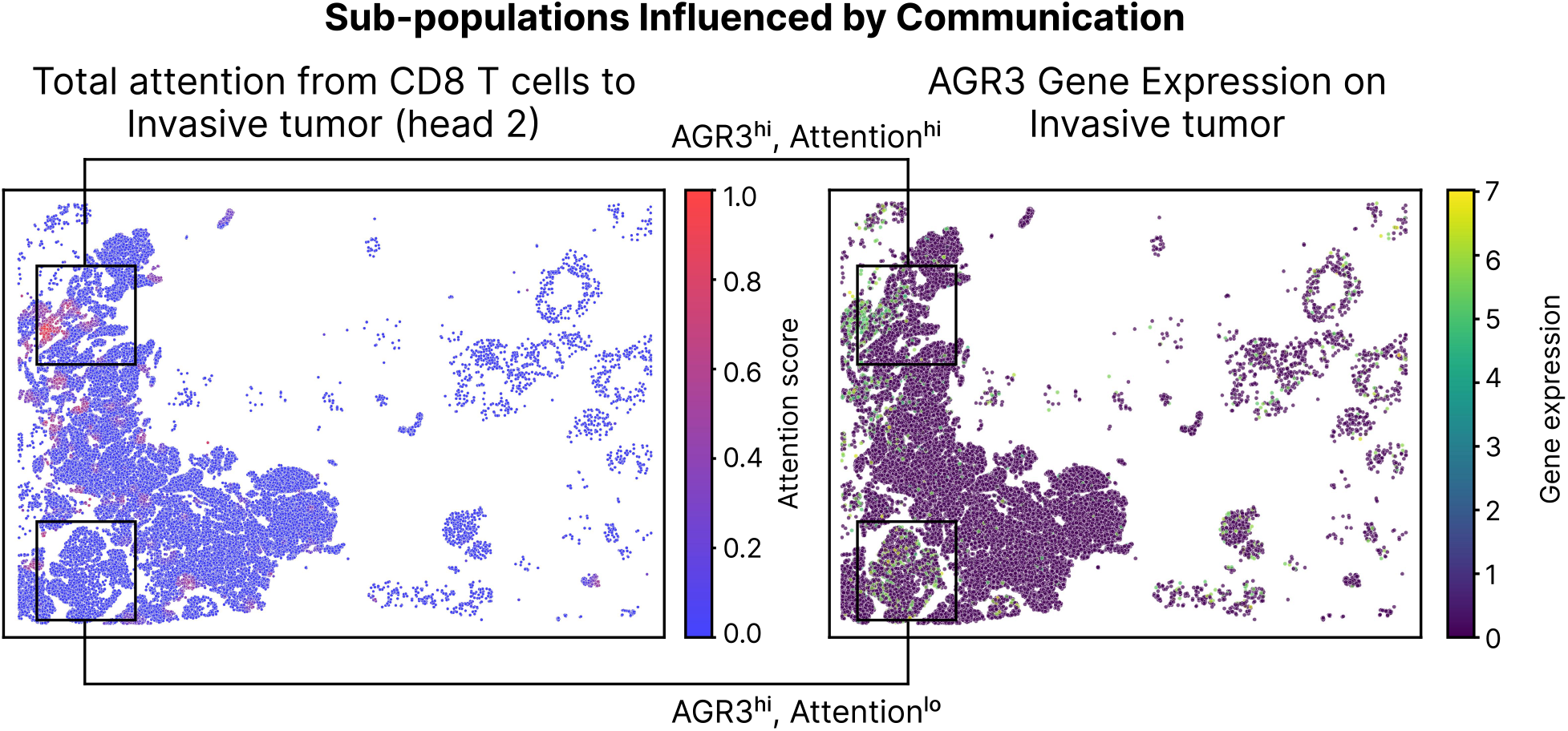
High attention from CD8^+^ T cells to invasive tumor overlaps with corresponding high *AGR3* expression at the tumor boundary as a result of interaction. Low attention overlapping with high *AGR3* expression – this high expression is as a result of potential segmentation from high DCIS 2 expression of the gene and the co-localization of these tumor cell types in that specific spatial region.

**Figure S7:**
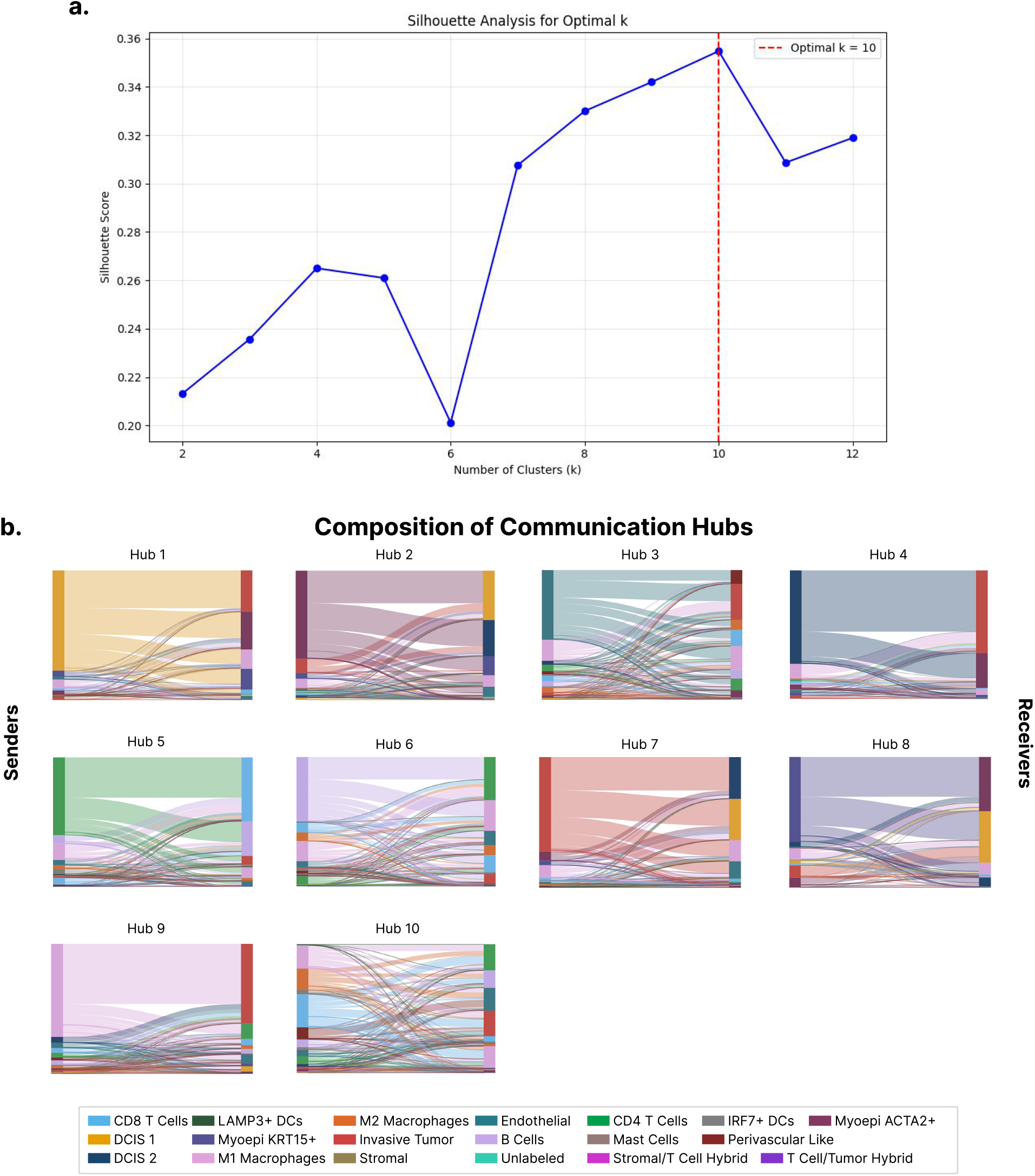
**a**. Plot displaying the silhouette score for cluster labels assigned by KMeans clustering of cells according to inferred AMICI interaction patterns, using various values of *k* to cluster, with the optimal value shown in the plot as *k* = 10. **b**. Compositions of sender and receivers in all communication hubs identified by AMICI.

**Figure S8:**
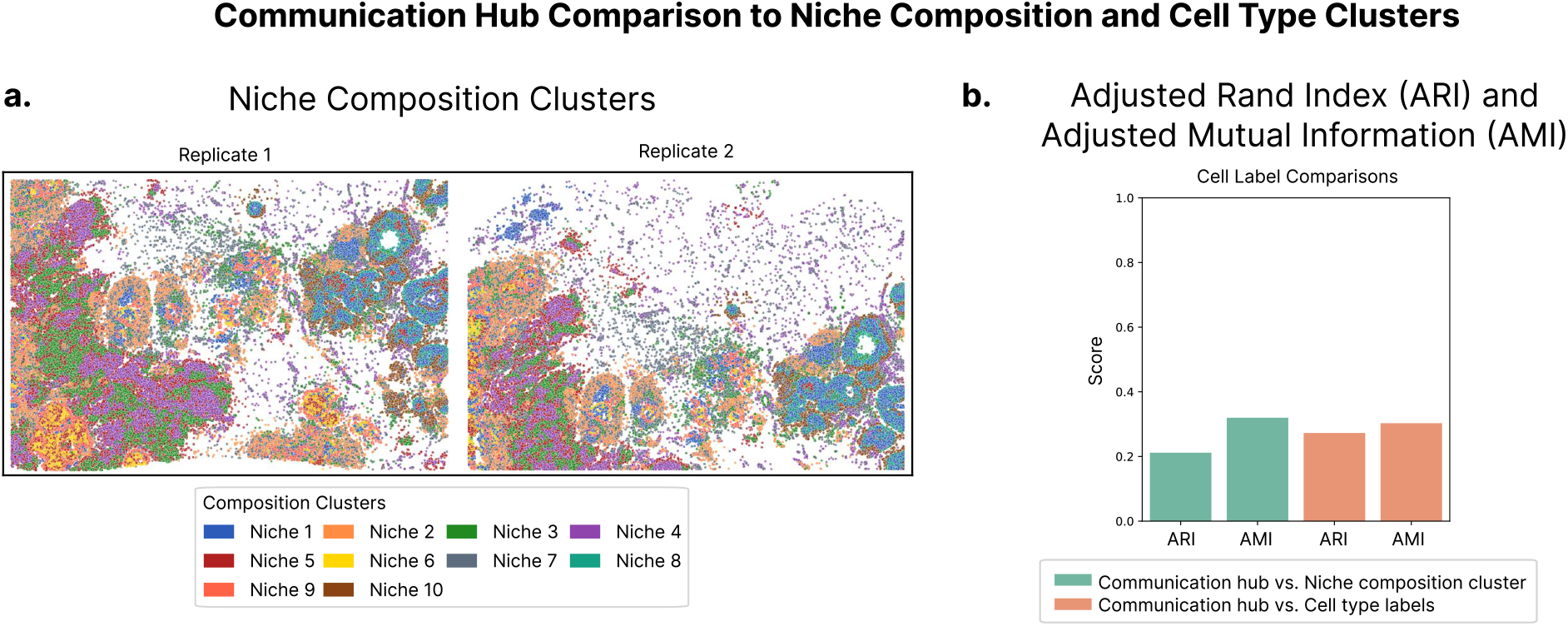
**a**. Spatial organization of tissue colored by niche composition clusters generated by the standard approach of clustering cells according to their neighbor cell type composition. **b**. Adjusted rand index (ARI) and adjusted mutual information (AMI) between the communication hubs identified by AMICI, and the niche composition clusters and cell type labels. The low similarity shows communication patterns are different across cell types and neighborhoods.

